# Sex dependence of opioid-mediated responses to subanesthetic ketamine

**DOI:** 10.1101/2022.09.06.506854

**Authors:** Tommaso Di Ianni, Matine M. Azadian, Sedona N. Ewbank, Michael Michaelides, Raag D. Airan

**Affiliations:** Department of Radiology, Stanford University School of Medicine, Stanford, CA 94305 USA; Biobehavioral Imaging and Molecular Neuropsychopharmacology Unit, National Institute on Drug Abuse Intramural Research Program, Baltimore, MD 21224 USA; Department of Psychiatry and Behavioral Sciences, Johns Hopkins University School of Medicine, Baltimore, MD 21205 USA; Department of Materials Science and Engineering, Stanford University School of Medicine, Stanford, CA 94305 USA; Department of Psychiatry and Behavioral Sciences, Stanford University School of Medicine, Stanford, CA 94305 USA

## Abstract

Subanesthetic ketamine rapidly and robustly reduces depressive symptoms in patients with treatment-resistant depression. While it is commonly classified as an *N*-methyl D-aspartate receptor (NMDAR) antagonist, our picture of ketamine’s mechanistic underpinnings is incomplete. Recent clinical evidence has indicated, controversially, that a component of the efficacy of ketamine in depression may be opioid dependent. Using pharmacological functional ultrasound imaging in rats, we found that blocking opioid receptors suppressed neurophysiologic changes evoked by ketamine, but not by a more selective NMDAR antagonist, in regions implicated in the pathophysiology of depression and in reward processing. Importantly, this opioid-dependent response was strongly sex dependent, as it was not evident in female subjects and was fully reversed by surgical removal of the male gonads. We observed similar opioid-mediated sex-dependent effects in ketamine-evoked structural plasticity and behavioral sensitization. Together, these results underscore the potential for ketamine to induce its affective responses via opioid signaling, and indicate that this opioid dependence may be strongly influenced by subject sex. These factors should be more directly assessed in future clinical trials.

**One-Sentence Summary:** Subanesthetic ketamine evokes opioid-mediated behavioral and neurophysiological effects in male, but not female, rats.

## Main Text

Major depression is a highly prevalent mood disorder and among the leading causes of disability worldwide (*1*). Traditional antidepressants are ineffective in the approximately 1/3 of patients with depression who develop treatment resistance (*2*). A single sub-anesthetic dose of (*R,S*)-ketamine rapidly and robustly reduces depressive symptoms (*3*, *4*), and a formulation of the (*S*)-ketamine stereoisomer is now FDA-approved for treatment-resistant depression (*5*). (*R*)-ketamine has also shown promising therapeutic potential in an open-label clinical trial (*6*).

The antidepressant action of ketamine is commonly attributed to its non-competitive antagonism at the glutamatergic *N*-methyl D-aspartate receptors (NMDAR) (*7*, *8*), but our picture of its underlying mechanisms is incomplete (*9*, *10*). Other experimental drugs acting via selective NMDAR antagonism or other downstream glutamate-mediated effects have provided minimal efficacy in clinical trials (*11*), while NMDAR-independent pathways have also been suggested to mediate the antidepressant action of ketamine (*12*). Recent clinical evidence suggests that pretreatment with naltrexone, a nonselective opioid receptor antagonist, dramatically attenuates the antidepressant effect of intravenous (i.v.) ketamine in humans (*13*, *14*). However, subsequent studies challenged these findings (*15*, *16*), warranting further investigation. Preclinical data also show diverging evidence: opioid receptor blockade suppressed the response to ketamine and its enantiomers in some studies (*17*–*19*), while others reported no significant effects (*20*). The potentially pivotal role of the opioid system in the antidepressant efficacy of ketamine raises concern for ketamine’s abuse liability and potential for dependence (*21*), as opioid signaling is thought to mediate the hedonic aspects of reward processing (*22*) and the reinforcing effects of drugs of abuse (*23*, *24*). This concern is particularly relevant in light of the ongoing opioid crisis (*25*). In addition, there has been relatively limited assessment of other biological variables, including subject sex, that may explain the heterogeneity of clinical and preclinical findings of the potential opioid dependence of ketamine’s therapeutic efficacy and adverse effects.

To determine how opioid receptor blockade affects ketamine-evoked neural activity changes, we assessed male and female rats using functional ultrasound imaging (fUSI). This neuroimaging modality is based on neurovascular coupling and closely tracks neural activity by way of high-resolution whole-brain maps of cerebral blood volume (CBV) (*26*, *27*). We first sought to establish that fUSI could resolve the acute effects of ketamine administration. We prepared male rats with a surgical craniotomy to allow ultrasound penetration and implanted a chronic polymeric prosthesis to enable repeated imaging (Fig. 1A). With the animals awake and restrained, we continuously recorded fUSI images at 2.5 mm rostral and 3.5 mm caudal to bregma (Fig. 1B). Ketamine (10 mg/kg, i.v.) evoked a rapid and sustained increase (peak at 3-5 min; 50 min duration) in CBV signal that extended over cortical and subcortical regions (Fig. 1C and Movie S1), in agreement with the reported excitatory action of low-dose NMDAR antagonism (*28*, *29*). We then infused varied doses of ketamine (0, 1, 5, and 10 mg/kg, i.v.) to determine the dose-response relationship of the CBV changes. We segmented the CBV signals in regions of interest (ROIs) obtained by registering the relevant slices from the Paxinos rat brain atlas onto a power Doppler vascular template and calculated the mean regional CBV time traces (Fig. 1D, S1). The traces presented a clear dose-response relationship, confirmed by statistical comparisons of the peak CBV (Fig. 1E) and area under the curve (AUC; Fig. S1B). Importantly, neither the peak CBV nor AUC were correlated to the local vascularization level, as measured by the regional baseline power Doppler signal, suggesting that the recorded changes were independent of the intrinsic vascular anatomy (Fig. S1C). Altogether, our results show that fUSI is able to image the neural effects evoked by acute ketamine infusions with a dose-dependent response, and confirm previous findings using different imaging modalities (*19*, *30*).

**Figure 1:**
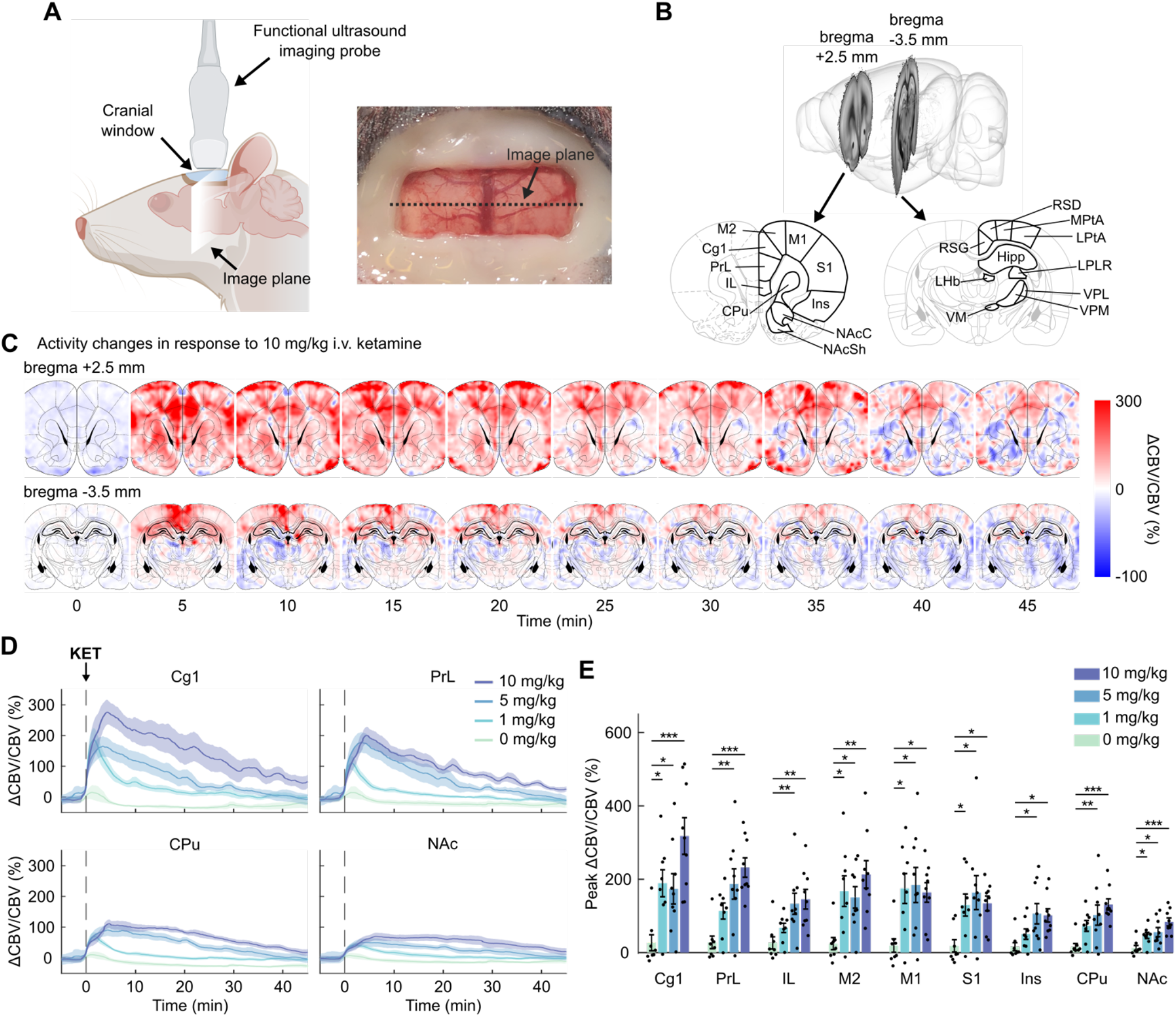
Functional ultrasound imaging of intravenous ketamine administration. (**A**) Schematic representation of the imaging setup. A surgical craniotomy enables ultrasound penetration, and a chronic prosthesis allows imaging over repeated sessions. (**B**) Coronal slices of the rat brain were imaged at bregma +2.5 mm and bregma −3.5 mm. The segmented regions of interest (ROIs) are highlighted on the relevant rat brain atlas. (**C**) Sequence of cerebral blood volume (CBV) coronal maps at bregma +2.5 mm and bregma −3.5 mm following administration of 10 mg/kg i.v. ketamine. The pixel intensity shows the CBV signals as a normalized difference with a pre-injection baseline (10 min). The time axis was zeroed at the time of ketamine injection. (**D**) The coronal maps were segmented and the CBV signals were averaged in the relevant ROIs. The plots show representative CBV time series in response to increasing doses of i.v. ketamine. Solid lines are mean and shaded areas are s.e.m. from *n* = 9/group (10 and 5 mg/kg) or *n* = 8/group (1 and 0 mg/kg). (**E**) Peak CBV in the segmented ROIs. Two-way mixed-effects ANOVA; within-subjects factor of region, *F*_3.08,92.55_ = 17.82, *P* < 0.0001; between-subjects factor of dose, *F*_3,30_ = 8.26, *P* = 0.0004; interaction, *F*_9.25,92.55_ = 3.38, *P* = 0.001. Two-tailed unpaired *t-*test, \**P* < 0.05; \*\**P* < 0.01; \*\*\**P* < 0.001. *n* = 8/9 rats per group. Data presented as mean ± s.e.m. KET: ketamine. Cg1: cingulate area 1. PrL: prelimbic cortex. IL: infralimbic cortex. M2: secondary motor cortex. M1: primary motor cortex. S1: primary somatosensory cortex. Ins: insular cortex. CPu: caudate putamen. NAcC: nucleus accumbens core. NAcSh: nucleus accumbens shell. LPtA: lateral parietal association cortex. MPtA: medial parietal association cortex. RSG: granular retrosplenial cortex. RSD: dysgranular retrosplenial cortex. Hipp: hippocampus. LHb: lateral habenula. LPLR: lateral posterior thalamic nucleus. VM: ventromedial thalamic nucleus. VPM: ventral posteromedial thalamic nucleus. VPL: ventral posterolateral thalamic nucleus.

To determine the presence of opioid-mediated effects, we pretreated two groups of male and female rats with subcutaneous (s.c.) injections of either vehicle (VEH; saline) or naltrexone (NTX; 10 mg/kg) followed by i.v. ketamine (KET; 10 mg/kg) or saline after 10 min (Fig. 2A). This ketamine dose reliably produces antidepressant-like effects in rat behavioral models (*12*). The 10 mg/kg naltrexone dose yields near complete mu opioid receptor occupancy in the mouse brain (*31*) and blocked the effect of (*S*)-ketamine on acute locomotion (*19*). Each rat was imaged three times under the treatment conditions of VEH+KET, NTX+KET, and NTX+VEH in a three-arm crossover design. Treatment conditions were assigned in randomized order to control for possible effects of prior drug exposure, and we allowed for a 7-day washout period between ketamine injections for full drug clearance.

**Figure 2:**
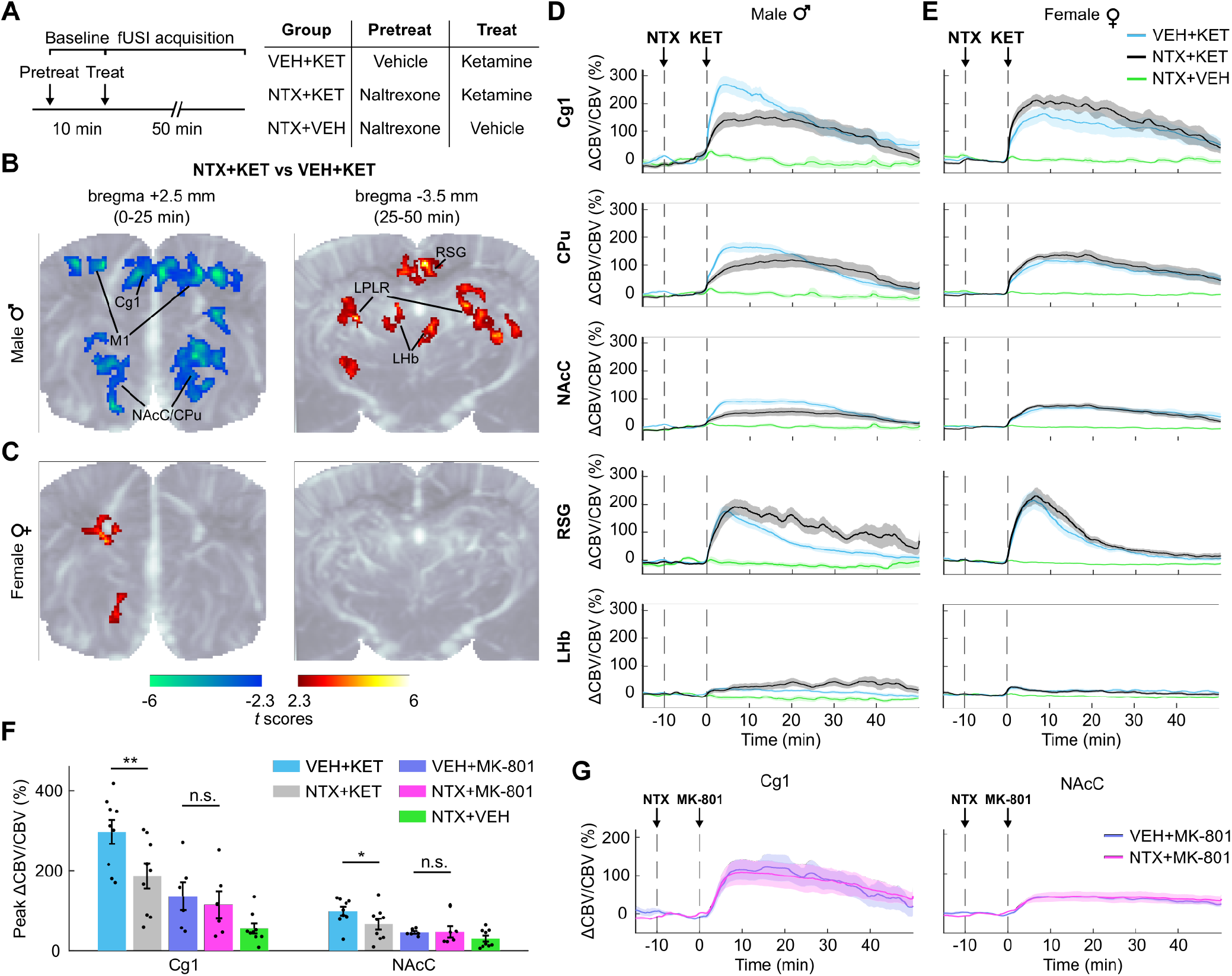
Pharmaco-functional ultrasound imaging reveals a sex-dependence of opioid-mediated effects of ketamine administration. (**A**) Rats received an s.c. injection of either vehicle (VEH; saline) or naltrexone (NTX; 10 mg/kg) followed by ketamine (KET; 10 mg/kg, i.v.) or vehicle after 10 min. Each animal was imaged three times under the treatment conditions of VEH+KET, NTX+KET, and NTX+VEH. (**B**, **C**) Functional maps in male and female rats. The *t* scores were calculated by contrasting the pixel-wise peak cerebral blood volume (CBV) in the NTX+KET versus VEH+KET groups. Statistically significant clusters are displayed overlaid on a power Doppler template (one cohort of *n* = 9 females and *n* = 9 males imaged at bregma +2.5 mm; one cohort of *n* = 9 females and *n* = 9 males imaged at bregma −3.5 mm; two-tailed paired *t-*test, *P* < 0.05). In male rats, functional maps show that naltrexone reduced peak activity in M1, Cg1, NAc core (NAcC), and CPu, and increased activity in RSG, LHb, and LPRL. There were only minor and less significant clusters in female rats. These results were also evident in the CBV time series (**D**, **E**). (**F**) A different cohort of male rats received an i.v. dose of 0.1 mg/kg MK-801 with naltrexone or vehicle pretreatment. The bar plot displays the peak CBV in the Cg1 and NAcC regions. Two-tailed paired *t-*tests (NTX+KET vs SAL+KET; NTX+MK-801 vs SAL+MK-801), \**P* < 0.05; \*\**P* < 0.01; n.s. not significant. Hedge’s *g* effect sizes: Cg1 = −1.20 (KET), −0.3 (MK-801); NAcC = −0.97 (KET), 0.04 (MK-801). *n* = 6. (**G**) CBV time series in Cg1 and NAcC in male rats receiving MK-801. *n* = 6. Data presented as mean ± s.e.m.

Functional maps contrasting the NTX+KET and VEH+KET groups revealed region-specific effects of naltrexone pretreatment (Fig. 2B, C; *P* < 0.05). Specifically, naltrexone pretreatment decreased ketamine-induced activity in the cingulate area 1 (Cg1) of the medial prefrontal cortex (mPFC), primary and secondary motor cortices (M1/2), dorsal striatum (CPu), and nucleus accumbens core (NAcC), and increased activity in the retrosplenial granular cortex (RSG), lateral habenula (LHb), and lateral posterior thalamic nucleus (LPLR) (Movie S2). Interestingly, these effects were only present in male rats, whereas females showed only minor and less significant differences with versus without naltrexone pretreatment (Fig. 3A). These results were also evident in the regional CBV time traces (Fig. 2D, E). The temporal dynamics of the naltrexone pretreatment effect were transient and region-dependent. We observed a biphasic effect where group differences in Cg1, M1/2, CPu, and NAcC occurred mostly in the 0-25 min interval, whereas the effect was delayed in the RSG, LHb, and LPLR (Fig. S2, S3). Importantly, naltrexone pretreatment produced no significant differences in CBV changes evoked by MK-801 (0.1 mg/kg; i.v.), a selective NMDAR antagonist (Fig. 2F, G; Fig. S4), suggesting that these opioid-dependent effects are specific to ketamine and independent of NMDAR action.

**Figure 3:**
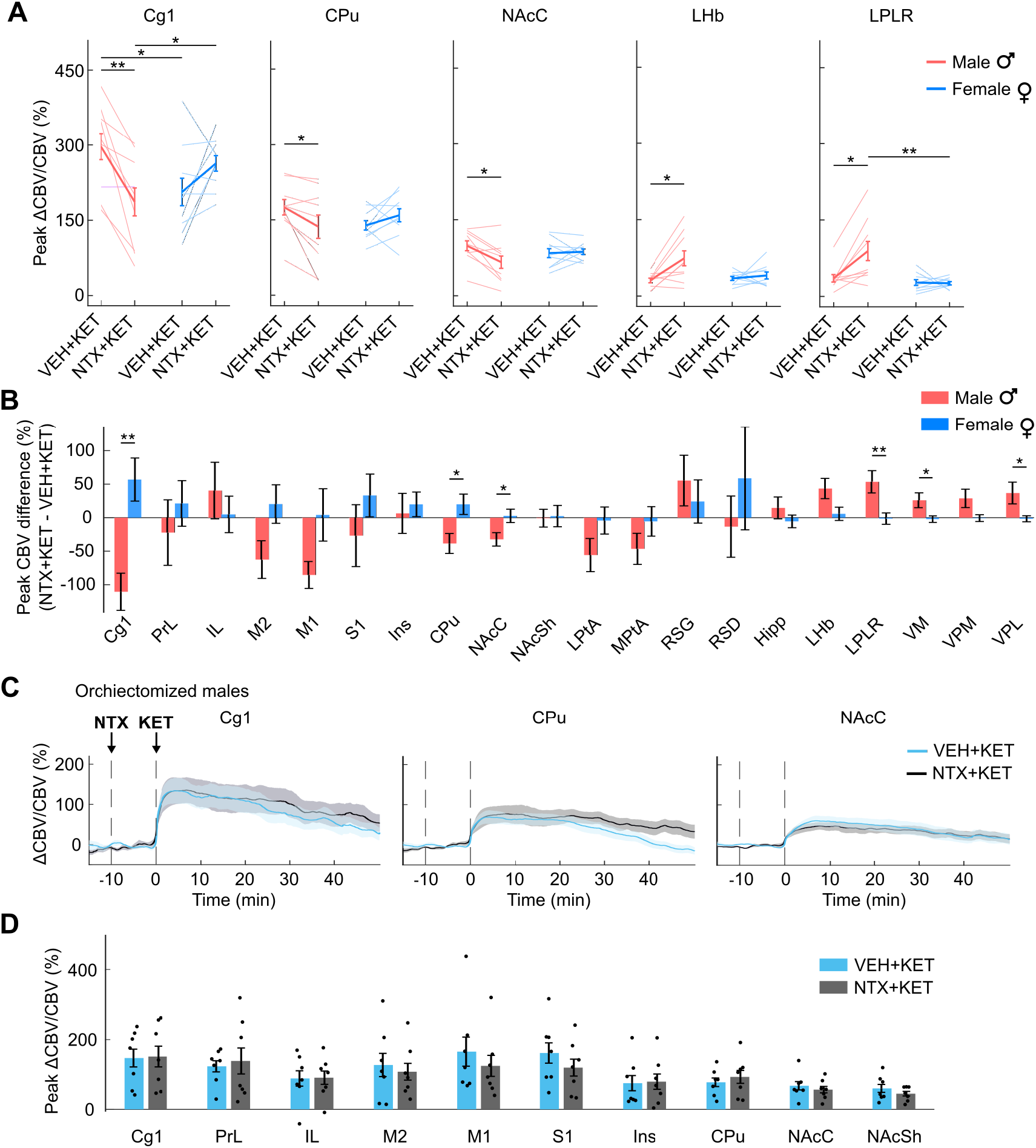
Sex-dependence of opioid-mediated effects of subanesthetic ketamine are region-specific. (**A**) Peak cerebral blood volume (CBV) changes in individual male and female rats receiving vehicle or naltrexone pretreatment. Two-tailed paired *t*-test (NTX+KET vs VEH+KET) or two-tailed unpaired *t*-test (F vs M), \**P* < 0.05; \*\**P* < 0.01. Hedge’s *g* effect sizes (NTX+KET vs VEH+KET): Cg1 = − 1.20 (M), 0.53 (F); CPu = −0.78 (M), 0.40 (F); NAcC = −0.97 (M), 0.08 (F); LHb = 0.87 (M), 0.17 (F); LPRL = 0.97 (M), −0.05 (F). Hedge’s *g* effect sizes (F vs M): Cg1 = −0.96 (VEH+KET), 0.97 (NTX+KET); LPLR: −1.34 (NTX+KET). *n* = 9 male; *n* = 9 female. (**B**) Peak CBV differences between the NTX+KET and VEH+KET treatments in individual rats were compared between males and females. One-way ANOVA for sex factor, *F*_1,358_ = 5.87, *P* = 0.016. Two-tailed unpaired *t*-test, \**P* < 0.05; \*\**P* < 0.01. Hedge’s *g* effect sizes: Cg1 = 1.77; CPu = 1.24; NAcC = 1.11; LPLR = −1.32; VM = −1.04; VPL = −1.02. Data presented as mean ± s.e.m. (**C**) A different cohort of male rats received a surgical orchiectomy. CBV time series show that the removal of gonadal hormones completely blocked the opioid-mediated effect of subanesthetic ketamine. *n* = 7. (**D**) Peak CBV in the segmented ROIs. Two-way ANOVA; within-subjects factor of region, *F*_9,54_ = 3.71, *P* = 0.0009; within-subjects factor of treatment, *F*_1,6_ = 0.31, *P* = 0.6; interaction: *F*_9,54_ = 1.33, *P* = 0.25. *n* = 7. Data presented as mean ± s.e.m.

To further investigate the effect of sex, we analyzed peak CBV differences between treatments in individual rats (Fig. 3B). We observed a significant effect of sex (one-way ANOVA, *F*_1,358_ = 5.87, *P* = 0.016), with specific differences between male and female rats in Cg1, CPu, NAcC, LPLR, VM, and ventroposterolateral thalamic nucleus (VPL) (post-hoc two-tailed unpaired *t*-test, *P* < 0.05). In a different cohort of male rats, we performed an orchiectomy (surgical removal of the gonads) to determine whether this sex-dependent effect was driven by endocrine factors rather than developmental sexual dimorphism of the brain (*32*). Interestingly, the effect of naltrexone pretreatment was completely blocked in orchiectomized males (Fig. 3C, D), suggesting a gating action of testosterone on the opioid-mediated response to ketamine. Collectively, our fUSI findings indicate that opioid receptors mediate the response to subanesthetic ketamine in key brain regions implicated in the pathophysiology of depression and in the processing of reward, and that this opioid-dependent effect is gated by the presence of male sex hormones.

Next, we sought to determine if the sex-dependent effects observed in our acute fUSI findings were reflected in physiological changes at the synaptic level. To this end, we used immunohistochemistry labeling of the postsynaptic density protein PSD-95 in fixed brain slices (Fig. 4A), as an indicator of ketamine-induced cellular structural plasticity. Synapse loss in prefrontal cortical neurons has been identified as a putative neurobiological substrate of depression, and ketamine reverses such synaptic deficits by restoring post-synaptic protein expression and functional spine density (*33*–*35*). Male and female rats received an antidepressant-like dose of 10 mg/kg ketamine intraperitoneally (i.p.), preceded by naltrexone (10 mg/kg, s.c.) or saline, and were perfused 24 h post infusion. Subanesthetic ketamine increased the expression of PSD-95 in the mPFC of both male and female rats compared to the saline-injected controls. In agreement with our imaging findings, naltrexone pretreatment blocked this effect in male (*P* = 0.0024) but not in female rats (*P* = 0.153) (Fig. 4B, C). Overall, these results indicate that the physiological changes at the synaptic level associated with ketamine’s antidepressant-like effect are indeed mediated by opioid receptors, but only in males.

**Figure 4:**
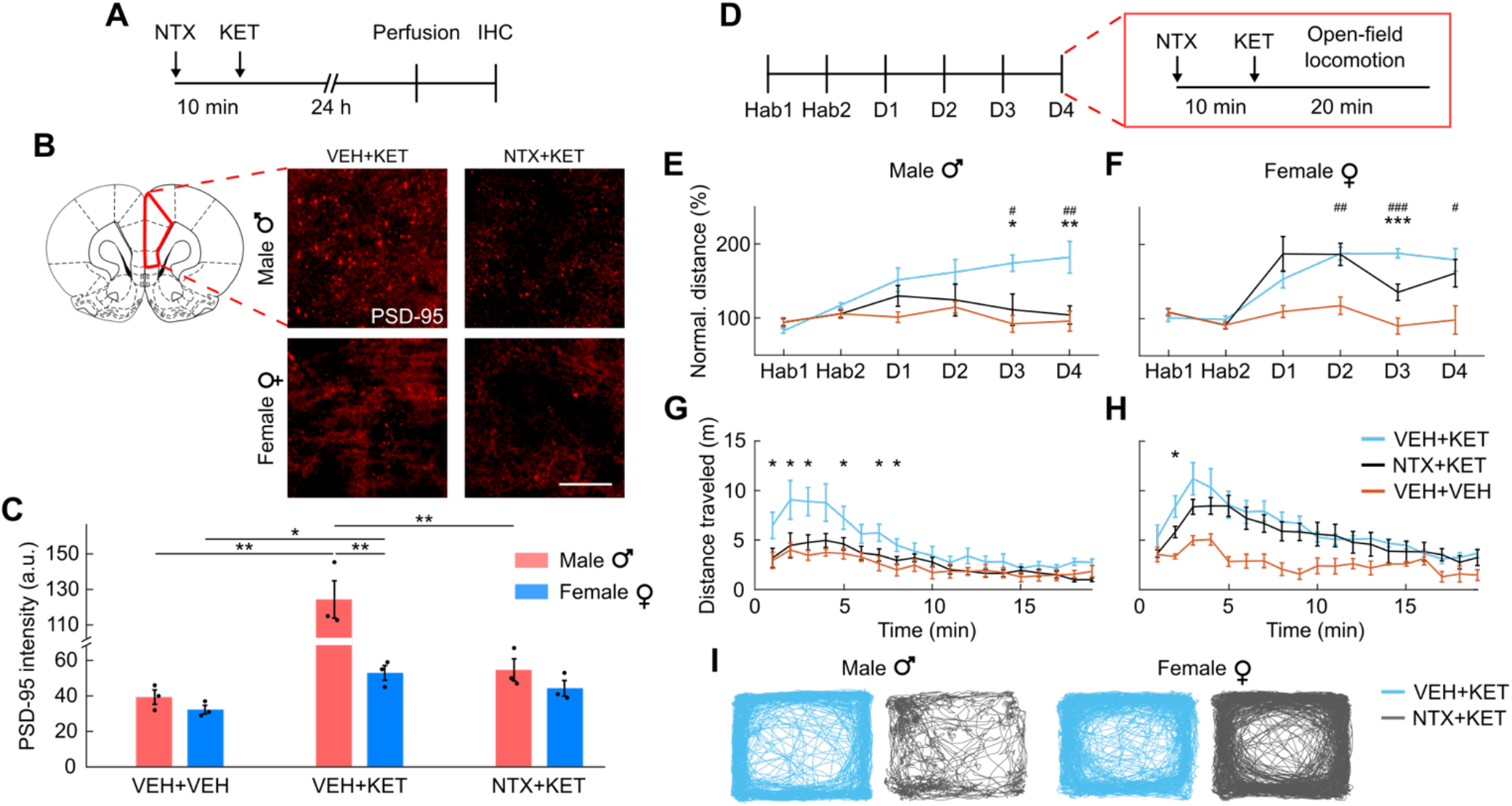
Sex-dependence of opioid-mediated structural plasticity and behavioral effects of subanesthetic ketamine. (**A**) Rats received an s.c. injection of naltrexone (NTX; 10 mg/kg) or vehicle (VEH) followed by ketamine (KET; 10 mg/kg, i.p.) or vehicle after 10 min and were perfused 24 h post ketamine infusion. Brain slices were analyzed by immunohistochemistry (IHC). (**B**) Representative PSD-95 stains in the rat prefrontal cortex at 20x magnification. Scale bar: 100 μm. (**C**) Quantification of PSD-95 intensity in the prefrontal cortex. Two-way ANOVA; treatment, *F*_2,12_ = 31.17, *P* < 0.0001; sex, *F*_1,12_ = 27.15, *P* = 0.0002; interaction, *F*_2,12_ = 13.59, *P* = 0.0008. Two-tailed unpaired *t*-test, \**P* < 0.05; \*\**P* < 0.01. Hedge’s *g* effect sizes: VEH+KET vs VEH+VEH = 3.84 (M), 2.81 (F); NTX+KET vs VEH+KET = −2.99 (M). *n* = 3/group. (**D**) Locomotor activity was measured for 20 min in an open-field arena. Male and female rats were pretreated with NTX or VEH followed by an injection of ketamine (10 mg/kg; i.p.) 10 min later on each of four days. Rats in the VEH+VEH group received two injections of saline. All animals were habituated for 2 days (Hab1-2) with vehicle only, followed by 4 daily sessions with ketamine with or without naltrexone pretreatment (D1-4). (**E**) Total distance traveled by male rats normalized to the mean of Hab1-2. Two-way mixed-effects ANOVA; between-subjects factor of treatment, *F*_2,17_ = 5.24, *P* = 0.017; within-subjects factor of session, *F*_2.74,46.6_ = 5.25, *P* = 0.004; interaction, *F*_5.48,46.6_ = 3.40, *P* = 0.009. VEH+KET vs VEH+VEH: two-tailed unpaired *t*-test, ^#^*P* < 0.05; ^##^*P* < 0.01; Hedge’s *g* effect sizes: D3 = 2.79; D4 = 1.70. NTX+KET vs VEH+KET: two-tailed unpaired *t*-test, \**P* < 0.05; \*\**P* < 0.01; Hedge’s *g* effect sizes: D3 = −1.25; D4 = −1.49. *n* = 8/group for VEH+KET and NTX+KET. *n* = 4 for VEH+VEH. (**F**) Total distance traveled by female rats. Two-way mixed-effects ANOVA; between-subjects factor of treatment, *F*_2,17_ = 9.89, *P* = 0.001; within-subjects factor of session, *F*_5,85_ = 15.1, *P* < 0.001; interaction, *F*_10,85_ = 4.16, *P* = 0.001. VEH+KET vs VEH+VEH: two-tailed unpaired *t*-test, ^#^*P* < 0.05; ^##^*P* < 0.01, ^###^*P* < 0.001; Hedge’s *g* effect sizes: D2 = 2.69; D3 = 4.65; D4 = 1.83. NTX+KET vs VEH+KET: two-tailed unpaired *t*-test, \*\*\**P* < 0.001; Hedge’s *g* effect sizes: D2 = −0.03; D3 = −1.97; D4 = −0.35. *n* = 8/group for VEH+KET and NTX+KET. *n* = 4 for VEH+VEH. (**G**, **H**) Distance traveled by male and female rats as a function of the time on day 4. Two-tailed unpaired *t*-test, VEH+KET vs NTX+KET, \**P* < 0.05, \*\*\**P* < 0.001. (**I**) Representative traces of body position during the open-field session at D4 in male and female rats. Data presented as mean ± s.e.m.

The antidepressant effect of a single ketamine infusion typically dissipates within hours. To achieve sustained remission, ketamine therapy requires repeated administration over the course of several weeks (*36*). Repeated exposure to drugs of abuse induces a progressive increase in locomotor behavior (i.e., locomotor sensitization) in rodents reflective of neuroadaptations of the mesolimbic dopamine system (*37*). As we found altered signaling in mesolimbic structures in our fUSI experiments, we aimed to investigate sex-dependent opioid-mediated effects on behavior in the context of repeated ketamine dosing. In a chronic open-field locomotor assay, we pretreated male and female rats with naltrexone (10 mg/kg, s.c.) or saline 10 min before injecting an antidepressant-like dose of ketamine (10 mg/kg, i.p.). Rats were habituated for 2 days with saline only, followed by 4 daily sessions with ketamine with or without naltrexone pretreatment (Fig. 4D). Repeated ketamine administration induced locomotor sensitization in both male and female rats (Fig. 4E-I). Importantly, naltrexone pretreatment completely blocked this effect in male rats, whereas in females there was a significant reduction of locomotor activity at day 3, but the effect was fully reversed at day 4 (Fig. 4H). Statistical analyses stratified by treatment showed a significant effect of sex (two-way mixed-effects ANOVA with within-subjects factor of sex and between-subjects factor of session; *F*_1,14_ = 6.22, *P* = 0.026), session (*F*_5,70_ = 8.82, *P* < 0.0001), and interaction (*F*_5,70_ = 3.14, *P* = 0.013) in rats pretreated with naltrexone, and a significant effect of session (*F*_2.85,39.9_ = 29, *P* < 0.0001) in rats pretreated with saline. Animals in the control group presented no significant effects. In summary, our results indicate that ketamine produced locomotor sensitization in both male and female rats, and pretreatment with the opioid receptor antagonist naltrexone completely blocked this effect in male rats only.

## Discussion

Our findings indicate that opioid receptors mediate, at least partly, the neural activity changes elicited by a subanesthetic dose of (*R*,*S*)-ketamine. These effects were mainly observed in the mPFC, NAc, and LHb, neural structures implicated in processing of reward and cognitive functions relevant for the pathophysiology of depression (*38*–*41*). Our results are in agreement with previous reports of ketamine directly recruiting opioid receptors in the mPFC (*19*) and LHb (*17*) *ex vivo*. Opioid signaling in the mPFC is consistent with ketamine’s antinociceptive and analgesic action (*42*), as this region executes descending pain control via functional connections with the periaqueductal gray (*43*), and activation of opioid receptors decreases LHb activity (*44*). The observed opioid-mediated response to ketamine in the NAc, a critical site for reward processing (*45*, *46*) and closely linked to the anhedonic symptoms of depression (*39*), may originate either from direct opioidergic action at this site (*47*) or via downstream projections from the mPFC (*48*). In a recent study, ketamine was also found to induce dopamine transients in the NAc similar to those evoked by cocaine (*49*), although it was concluded to have limited addiction potential.

Interestingly, these opioid-mediated effects were markedly dependent on the presence of male sex hormones, as they were not evident in female subjects and were fully reversed by surgical removal of the male gonads. These acute observations were reflected in downstream changes in synaptic plasticity in the mPFC, a putative target of the antidepressant action of ketamine (*8*). Importantly, repeated dosing of subanesthetic ketamine induced behavioral sensitization, a robust measure of the neurobehavioral adaptations caused by repeated exposure to drugs with abuse liability (*50*), in both male and female rats, and blocking the opioid receptors also suppressed this effect, but only in male subjects.

Proper comparisons of ketamine’s therapeutic and adverse effects in male and female patients are lacking, possibly due to the limited statistical power for such subgroup analysis in current clinical trials, although sex differences have previously been reported at an anecdotal level (*51*). Sex-dependent ketamine and norketamine pharmacokinetics have been observed (*52*), and diverging correlations between treatment outcomes and depression-related inflammatory cytokines have been reported in male and female subjects (*53*). Moreover, preclinical investigations showed sex differences in the pharmacokinetics of ketamine and its metabolites, as well as in behavioral and physiological readouts (*12*, *52*, *54*). Our results represent a critical step forward in elucidating both sex-dependent and opioid-based mechanisms underlying the action of subanesthetic ketamine. The timeliness for such investigation is made more urgent by the current widespread administration of ketamine, in both its racemic and isomer-specific forms, in patients with treatment-resistant depression and other affective disorders. Indeed, it is possible that the current heterogeneity in the clinical and preclinical findings and the associated controversy surrounding the potential opioid dependence of ketamine’s clinical effects could reflect the existence of explanatory demographic-based biological variables, such as sex.

There are several limitations to this study. First, subanesthetic ketamine causes transient cardiovascular effects in both humans and rats (*5*, *55*), which may confound the interpretation of our fUSI results. However, the dynamics of ketamine’s activity and the effect of naltrexone pretreatment varied substantially between brain regions and throughout the scanning time, with opposing effects in different brain regions; therefore, we consider that the observed effects were unlikely to reflect a simple global cardiovascular modulation. In addition, previous studies showed similar patterns of neural activity with complementary modalities including pharmacological magnetic resonance imaging (*56*) and [^18^F]-fluorodeoxyglucose positron emission tomography (*19*). Second, in female rats we did not control for the estrous cycle at the time of injection (*57*). However, the physiological effect of ketamine is not dependent on the estrous phase (diestrus or proestrus) (*34*), and surgical ovariectomy did not alter the plasma levels of ketamine and its metabolites in mice (*52*). Moreover, our randomized study design should mitigate any confounds related to the estrous phase, as female rats in each treatment group were likely imaged during different phases of the estrous cycle.

In summary, our results establish that opioid blockade can modulate neural activity, cellular physiologic, and behavioral changes induced by ketamine, but only in male subjects. Therefore, it is imperative that future clinical trials focus on sex as a biological variable in assessing the affective responses to subanesthetic ketamine, including its antidepressant efficacy, especially with respect to potential abuse liability or withdrawal type responses upon discontinuation (*58*, *59*).

## Acknowledgements

We would like to thank Profs. Nolan Williams, Boris Heifets, and Robert Malenka for invaluable conversations, Profs. Jeremy Dahl, Kim Butts Pauly, and Katherine Ferrara for helpful advice on ultrasound methodologies, Dr. Keith Murphy for feedback on figure crafting, and all members of the Airan and Michaelides labs for helpful discussions.

## Funding

Seed Grant from the Stanford Wu Tsai Neurosciences Institute (RDA).

NIH BRAIN Initiative (NIH/NIMH RF1MH114252 to RDA).

NIH HEAL Initiative (NIH/NINDS UG3NS115637 to RDA).

NIDA Intramural Research Program (ZIA000069 to MM).

Stanford University School of Medicine Dean’s Postdoctoral Fellowship (TDI).

Ford Foundation Fellowship Program of the National Academies of Sciences, Engineering, and Medicine (MMA).

National Science Foundation Graduate Research Fellowship Program (SNE).

## Author Contributions

Designed and performed experiments, analyzed data, and wrote the manuscript: TDI.

Designed and performed experiments, analyzed data: MMA.

Designed experiments, contributed to writing the manuscript, funding acquisition, supervision: RDA.

Performed experiments: SNE.

Contributed to the design of experiments: MM.

All coauthors reviewed the manuscript and provided comments.

## Data and materials availability

Data and codes are available from the corresponding author upon reasonable request.

## Supplementary Materials

Materials and Methods

Figs. S1 to S5

Movies S1 to S2

