## Supplementary Materials for "Sex dependence of opioid-mediated responses to subanesthetic ketamine"

**This PDF file includes:**

Materials and Methods
Figs. S1 to S5
Captions for Movies S1 to S2

**Other Supplementary Materials for this manuscript include the following:**

Movies S1 to S2

### Materials and methods

#### Animals

The experimental protocol for all the animal procedures was approved by the Institutional Animal Care and Use Committee at Stanford University. Male and female Long Evans rats (Charles River Laboratories) were used in the experiments. All animals were 9-10 weeks old and weighed 278 $\pm$ 40 g (mean $\pm$ s.d.) when they entered the study. Animals had *ad libitum* access to water and food for the entire duration of the experimental protocols. Rats were housed in a temperature-controlled vivarium on a 12-h light-dark cycle (lights on at 7 AM; lights off at 7 PM) and were acclimated to their home cage for one week before experimentation. In case of surgical procedures, the animals were singly housed following the surgery.

#### Drugs

Naltrexone hydrochloride (Tocris Bioscience) was suspended in 0.9% sterile saline to obtain a 10 mg/mL solution. MK-801 (Tocris Bioscience) was suspended in 0.9% sterile saline to obtain a 0.1 mg/mL solution. Ketamine hydrochloride (Dechra Veterinary Products) was diluted in 0.9% sterile saline to obtain 1, 5, and 10 mg/ml solutions. All drugs were administered in a bolus injected volume of 1 mL/kg.

#### Surgical procedures

##### *Craniotomy*

Rats received a bilateral surgical craniotomy and chronic prosthesis implantation as previously reported (*60*). Briefly, animals were anesthetized with 3.5% isoflurane in 100% oxygen, and anesthesia was maintained with 1.5% isoflurane. The incision region was prepared by shaving the skin using a depilatory cream. Rats were then placed in a stereotaxic frame for head fixation and orientation. Body temperature was maintained at 37 °C by a warming pad with rectal probe monitoring (RightTemp Jr.; Kent Scientific). Heart rate and arterial oxygen saturation were monitored by a pulse oximeter (MouseStat Jr.; Kent Scientific). Anti-inflammatory (dexamethasone, 1 mg/kg; IP) was administered to prevent brain swelling and inflammation. The incision site was disinfected by applying alternating povidone-iodine and 75% EtOH, and a skin incision was performed. The bone was cleaned with 75% EtOH and a window (5 mm AP × 10 mm ML centered at bregma +2.5 mm) was marked on the skull with a surgical pen. The bone around the window was pretreated using a bonding agent (iBOND Total Etch; Kulzer). We then cut parietal and frontal bone fragments using a handheld high-speed drill with a 0.7 mm drill bit (Fine Science Tools). We gently removed the bone flaps paying attention to avoid damaging the dura mater, and sealed with dental cement (Tetric EvoFlow; Ivoclar Vivadent) a 125-μm polymethylpentene film covering the cranial window. The space between the dura and the prosthesis was filled with 0.9% sterile saline. A dose of 0.5 mg/kg Buprenex SR was administered subcutaneously for analgesia. The animals were allowed to recover for 1 week before the first imaging session.

*Orchiectomy*

To assess the effect of sex hormones, adult male rats were orchiectomized as previously reported (*61*). The animals were anesthetized with 3.5% isoflurane in 100% oxygen, and anesthesia was maintained with 1.5% isoflurane. The incision site was prepared by shaving and disinfecting the skin as described above. An incision of about 10 mm was made on the ventral side of scrotum along the midline. The testicular content was exposed, the vas deferens and blood vessels were clamped to prevent bleeding, and the testicles were removed. The incision was then closed with monofilament sutures. We waited for 10 days before experimentation to allow for recovery and testosterone washout.

#### Pharmaco-functional ultrasound imaging

*Ultrasound system and power Doppler processing*

A Vantage 256 research scanner (Verasonics Inc.) was connected to a linear array transducer (Vermon; 128 elements, lateral pitch of 100 μm) operating at a 15-MHz center frequency. The imaging probe was housed in a custom 3-D printed holder mounted on a motorized positioning system. For acoustic coupling, we used ultrasound gel that was centrifuged to remove air bubbles. The imaging sequence consisted of five tilted plane waves (-6°, -3°, 0°, 3°, 6°) emitted with a pulse repetition frequency of 19 kHz. Two plane waves were averaged for each angle to increase the signal-to-noise ratio. We acquired data for 200 compound frames at a rate of 1 kHz, and the frames were beamformed in a regular grid of pixels with in-plane resolution of 100 μm × 100 μm. Beamforming was performed in real-time in an NVIDIA Titan RTX using a GPU beamformer (*62*).

Sequences of 200 compound ultrasound frames were processed offline in MATLAB (MathWorks, Inc.) for clutter filtration and power Doppler computation. To eliminate the Doppler signal component originating from the stationary tissue, we used a $5^{th}$-order temporal high-pass Butterworth filter with a cutoff frequency of 40 Hz and a singular value decomposition filter that eliminates the first singular value (*63*). The power Doppler intensity at each pixel was calculated by squaring and averaging the filtered Doppler signals. The final power Doppler frame rate was 1 frame/sec.

*Imaging session*

At the beginning of each imaging session, rats were briefly anesthetized with isoflurane and a catheter was placed in the tail vein for vascular access. While under anesthesia, animals were placed in a plastic restraint cone (Stoelting Co.) and positioned in a custom head-restraining apparatus (*64*). Oxygen was flowed through the nose cone to prevent hypoxia. The ultrasound probe was positioned over the slice of interest. The relevant brain atlas slice was plotted overlaid on the real-time power Doppler images to facilitate accurate probe positioning based on vascular landmarks. With the animal in the imaging apparatus, we waited for 30-45 min before data acquisition to allow for complete isoflurane clearance. An s.c. injection of naltrexone or vehicle was performed, followed by an i.v. injection of drug (ketamine or MK-801) or vehicle after 10 min. After the i.v. injection, the catheter was flushed with 200 μL of sterile saline. We acquired data continuously for up to 50 min following drug administration.

*Functional ultrasound data pre-processing*

To prevent motion artifacts in the processed CBV signals, translational and rotational movements were corrected by applying a motion correction algorithm to the image time series. For each acquisition, a power Doppler template was calculated via median filtering of the first 500 images. Then, all power Doppler frames in the same acquisition were registered to the template using a rigid transformation that included rotations, translations, and cubic interpolations. A filter was used to remove registered data frames affected by excessive motion or other artifacts. This filter was adapted from previously published code (*65*). Each power Doppler dataset was then manually registered to the relevant slice of the Paxinos brain atlas (*66*) (at bregma +2.5 mm or bregma -3.5 mm).

*Cerebrovascular time traces*

The pixel-wise relative CBV signal was calculated as the normalized difference with a baseline (i.e., $\Delta CBV/CBV = (CBV_t - CBV_0)/ CBV_0$). For each acquisition, the baseline was calculated by averaging 10 min of power Doppler data immediately before drug administration. The regional time traces were computed by spatially averaging the pixel ΔCBV/CBV signals in the relevant segmented ROIs in each brain slice (Fig. 1a).

*Functional maps*

To assess the effect of ketamine administration and naltrexone pretreatment, we used an approach similar to direct pharmaco-fMRI (*56*), where we used pixel-wise statistical inference to analyze group-level differences in peak CBV signal. For each pre-processed power Doppler acquisition, an image was created by calculating the temporal CBV peak at each spatial location. Peak CBV images were registered to the template atlas space by performing a rigid transformation, and *t* scores were calculated for the contrasted groups (NTX+KET vs VEH+KET; two-tailed paired *t* test). Thresholded *t* scores were corrected for multiple comparisons across each slice using a cluster-size threshold of 34 contiguous pixels. The threshold was determined via Monte Carlo simulations using the 3dClustSim program of the AFNI library (*67*) to obtain an overall cluster $P < 0.05$, family-wise error rate corrected. Color-coded functional maps were displayed overlaid on a power Doppler template to enable a visual comparison of the analyzed groups.

#### Postsynaptic density protein PSD-95

##### *Drug administration and immunohistochemistry*

Rats were administered an s.c. injection of 10 mg/kg naltrexone or vehicle. After 10 min, an i.p. injection of 10 mg/mg ketamine or vehicle was performed, and the animals were returned to their home cage. After 24 h post-ketamine, the animals were anesthetized with isoflurane (5%) and transcardially perfused with 1x phosphate-buffered saline (PBS) followed by 4% paraformaldehyde (PFA) diluted in PBS. Brains were extracted and fixed overnight in 4% PFA, then subsequently washed in PBS and frozen in embedding medium. Coronal sections 40 μm thickness were cut on a CM1800 Cryostat (Leica Microsystems), transferred to tissue storage solution (30% sucrose and 30% ethylene glycol in 0.1M PB), and stored at –20 °C until immunohistochemical processing. Four tissue sections per rat (from Bregma +2.7 to +1.2; one 40 μm section every 400 μm) were selected for PSD-95 and DAPI labeling. Floating sections were rinsed with PBS then blocked with 4% normal goat serum and 0.3% Triton-X 100 diluted in PBS. Sections were then incubated overnight at 4°C in primary antibody, rabbit monoclonal to PSD95 (ab238135; Abcam). Following incubation, sections were rinsed in PBS and incubated in secondary antibody, goat-anti-rabbit Alexa Fluor 555 (Invitrogen) at 1:500 for 2h. Sections were then mounted on super-plus glass slides (VWR), airdried in the dark, and cover-slipped with hard-set mounting medium containing DAPI (Vector Labs).

##### *Microscopy and image analysis*

Images for PSD95-stained sections were acquired on a Keyence BZ-X800 fluorescence microscope (Keyence Corp.). Acquisition settings remained strictly constant between all images acquired at the same magnification. Specific ROIs were chosen to sample the mPFC at approximately the infralimbic, prelimbic, and cingulate area 1. High-resolution z-stacks of each ROI were acquired using a 40x magnification with a step size of 0.4 μm and total depth of 6 μm. Each Z-stack image set was merged and analyzed with BZ-X Advanced Analysis Software (Keyence Corp.). Signals above thresholded background were used for manual ROI segmentation to calculate the area of mean fluorescent signal intensity of each ROI, averaged across the four sections collected per animal. Mean fluorescent intensity is reported in arbitrary units.

#### Locomotor sensitization

All behavioral tests were performed in an environmentally controlled room. Open-field locomotor activity was recorded in a custom-built white Plexiglas apparatus (90 cm × 90 cm × 40 cm) divided in four equal compartments. Videos were collected for batches of 4 animals using an overheard camera placed at the center of the field. Animals in each batch were randomized for sex and treatment group (SAL + KET; NTX + KET; SAL + SAL). Prior to the behavioral tests, rats were handled for 3 days to acclimate to the experimenter and reduce stress. Then, locomotor activity was recorded for a total of 6 days. In the first two habituation days (HAB1/2), rats received an s.c. injection of vehicle and were then returned to their home cage. After 10 min, rats received an i.p. injection of vehicle and were immediately placed at the center of the arena, where they were allowed to freely explore for 20 min while locomotor activity was recorded. In the following 4 days (D1/4), rats were administered an s.c. injection of vehicle or naltrexone (10 mg/kg) and returned to their home cage. After 10 min, rats received an i.p. injection of ketamine (10 mg/kg) and were placed at the center of the arena while locomotion was recorded. Animals in the control group (SAL + SAL) continued to receive vehicle injections for the entire duration of the experiment. The compartments were thoroughly cleaned with Virkon between each recording session to control for scent-related confounds. White noise (65 dB) was played during the sessions to attenuate any external noise. Both a male (TDI) and a female (SNE) experimenter conducted the tests to control for any confounds introduced by the experimenter's sex (*68*). All behavioral tests were performed at the end of the light cycle, between 4:00 PM and 7:00 PM. The videos were analyzed in ToxTrac (*69*) to track the instantaneous animal center position and quantify distance traveled. During habituation, female rats showed higher locomotion than males (Fig. S5A; two-sided unpaired $t$-test, $P = 0.0006$), therefore we normalized the distance traveled to the habituation baseline to isolate the effect of ketamine.

General statistical analysis

Rats were randomly assigned to treatment conditions. When within-subject factors were present in the ANOVA, Mauchly's test for sphericity was performed to determine whether the sphericity assumption was satisfied. In cases where the assumption was violated, we used a Greenhouse-Geisser adjustment to the degrees of freedom. Pairwise post-hoc comparisons were performed in case of significant ANOVA effects. In the pairwise tests, multiple comparisons were controlled using Benjamini–Hochberg false-discovery-rate (FDR) correction ($\alpha = 0.05$). All comparisons were two-tailed. We calculated effect sizes using Hedge's $g$. Statistical tests, sample sizes $n$, corrected $P$ values, and effect sizes $g$ are reported for each analysis in the text and figure captions. All statistical analyses were performed using custom scripts in R Studio and MATLAB.

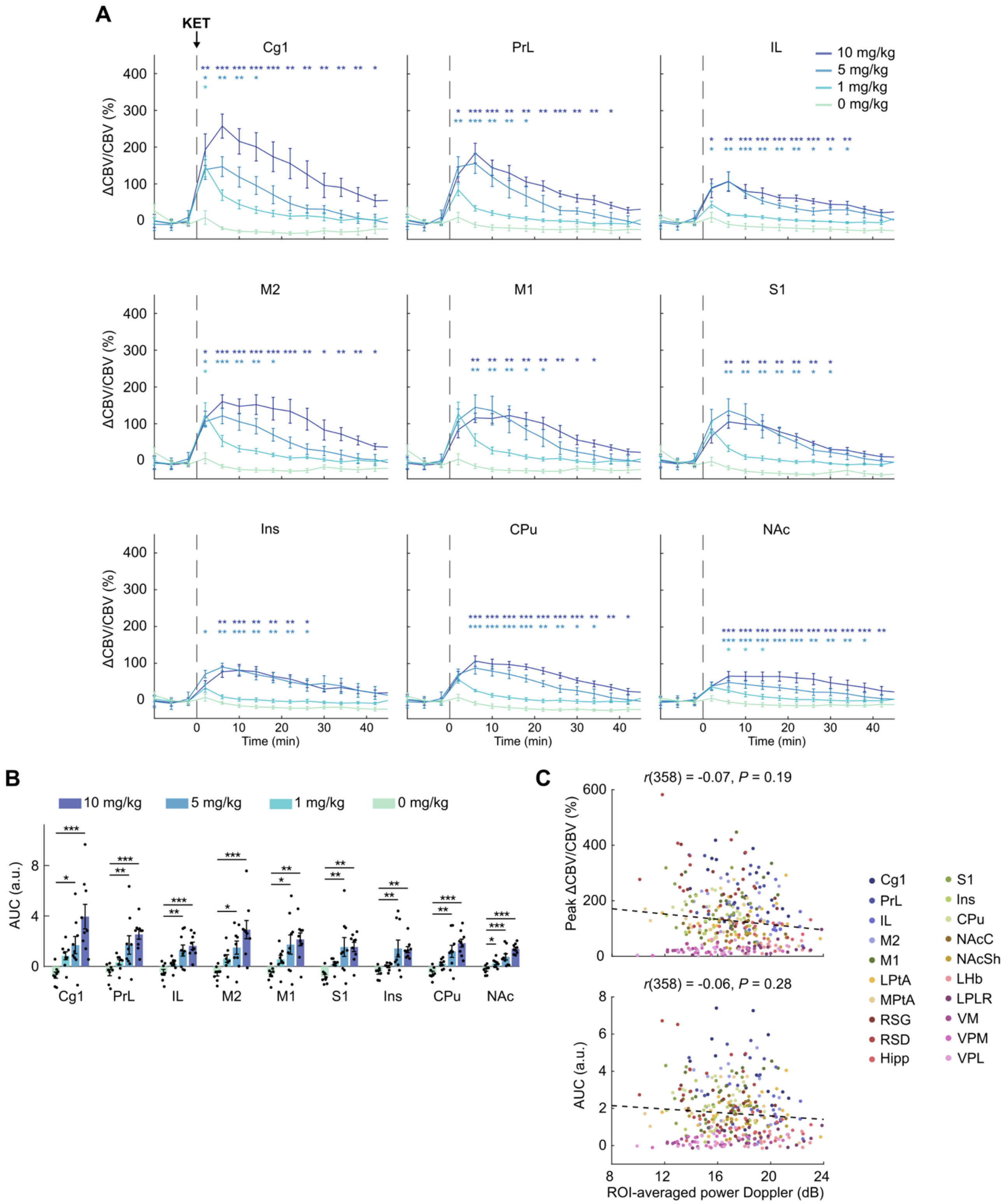


**Figure S1: Dose-dependent response of pharmacologic functional ultrasound imaging of ketamine induced responses.** (**A**) Regional cerebral blood volume (CBV) signals in the segmented regions of interest were averaged in 4-min intervals for statistical analysis. Lines are mean and error bars are s.e.m. from $n$ = 9/group (10 and 5 mg/kg) or $n$ = 8/group (1 and 0 mg/kg). Three-way mixed-effects

ANOVA; within-subjects factor of time, $F_{14,420} = 26.01$, $P < 0.0001$; within-subjects factor of region, $F_{8,240} = 5.11$, $P < 0.0001$; between-subjects factor of dose, $F_{3,30} = 9.9$, $P < 0.001$; interactions: dose × time, $F_{42,420} = 6.55$, $P < 0.0001$; dose × region, $F_{24,240} = 2.57$, $P < 0.001$; time × region, $F_{112,3360} = 6.95$, $P < 0.0001$; dose × time × region, $F_{336,3360} = 2.37$, $P < 0.0001$. Two-tailed unpaired *t*-test against the control group (0 mg/kg), *$P < 0.05$, **$P < 0.01$, ***$P < 0.001$. (**B**) Area under the curve (AUC) of the CBV signals in the segmented ROIs with intravenous ketamine at increasing doses (0, 1, 5, and 10 mg/kg). Two-way mixed-effects ANOVA; within-subjects factor of region, $F_{2.78,83.35} = 5.32$, $P = 0.003$; between-subjects factor of dose, $F_{3,30} = 10.47$, $P < 0.0001$; interaction, $F_{8.33,83.35} = 2.62$, $P = 0.012$. Two-tailed unpaired *t*-test, *$P < 0.05$; **$P < 0.01$; ***$P < 0.001$. $n$ = 8/9 rats per group. Data presented as mean ± s.e.m. (**C**) Correlation of ROI-averaged power Doppler with regional peak CBV and regional AUC. Spearman's rank correlation, 360 ROI-segmented values from $n$ = 36 animals.

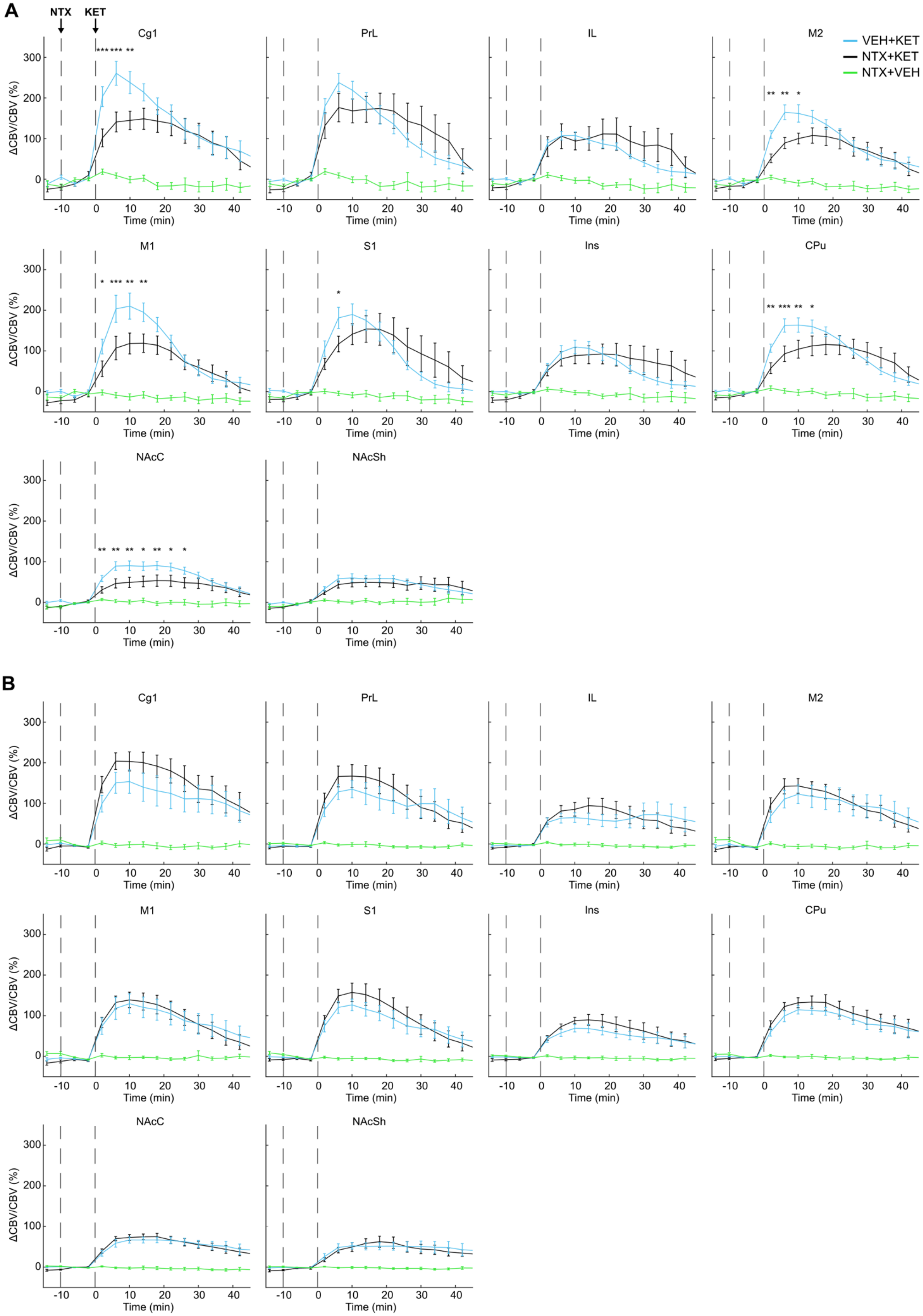

A
NTX
KET
Cg1
PrL
IL
M2
VEH+KET
NTX+KET
NTX+VEH
M1
S1
Ins
CPu
NAcC
NAcSh
ΔCBV/CBV (%)
Time (min)
B

**Figure S2: Regional cerebral blood volume (CBV) time series following ketamine administration at bregma +2.5 mm.** (**A, B**) The regional CBV signals in the segmented ROIs imaged at bregma +2.5 mm in male (**A**) and female (**B**) rats were averaged in 4-min intervals. Lines are mean and error bars are s.e.m. from $n = 9$ male rats and $n = 9$ female rats. Four-way mixed-effects ANOVA with the within-subjects factors of treatment, region, and time and the between-subjects factor of sex. Significant effects: treatment ($F_{2,32} = 46.05$, $P < 0.0001$), time ($F_{15,240} = 78.76$, $P < 0.0001$), region ($F_{9,144} = 18.23$, $P < 0.0001$), sex × time ($F_{15,240} = 2.41$, $P = 0.003$), treatment × time ($F_{30,480} = 19.96$, $P < 0.0001$), treatment × region ($F_{18,288} = 7.46$, $P < 0.0001$), time × region ($F_{135,2160} = 16.85$, $P < 0.0001$), sex × treatment × time ($F_{30,480} = 2.24$, $P = 0.0002$), sex × time × region ($F_{135,2160} = 1.37$, $P = 0.004$), treatment × time × region ($F_{270,4320} = 5.42$, $P < 0.0001$), sex × treatment × time × region ($F_{270,4320} = 1.33$, $P = 0.0004$). Two-tailed paired *t*-test, VEH+KET vs NTX+KET, *P < 0.05, **P < 0.01, ***P < 0.001.

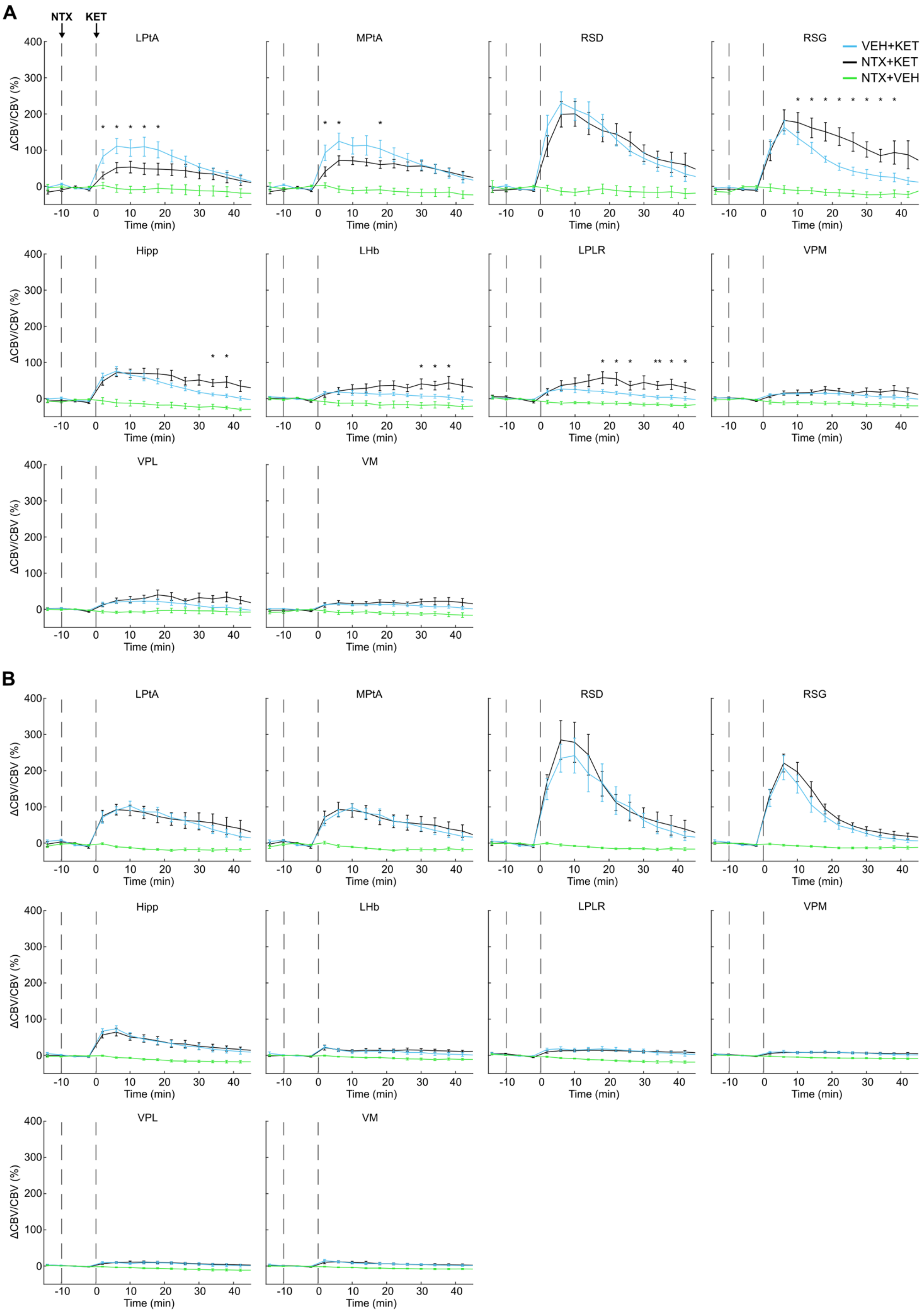
A
NTX
KET
LPtA
MPtA
RSD
RSG
VEH+KET
NTX+KET
NTX+VEH
ΔCBV/CBV (%)
Time (min)
Hipp
LHb
LPLR
VPM
VPL
VM
B
LPtA
MPtA
RSD
RSG
Hipp
LHb
LPLR
VPM
VPL
VM

227

**Figure S3: Regional cerebral blood volume (CBV) time series following ketamine administration at bregma -3.5 mm.** (**A, B**) The regional CBV signals in the segmented ROIs imaged at bregma +2.5 mm in male (**A**) and female (**B**) rats were averaged in 4-min intervals. Lines are mean and error bars are s.e.m. from $n$ = 9 male rats and $n$ = 9 female rats. Four-way mixed-effects ANOVA with the within-subjects factors of treatment, region, and time and the between-subjects factor of sex. Significant effects: treatment ($F_{2,32}$ = 74.72, $P$ < 0.0001), time ($F_{15,240}$ = 49.19, $P$ < 0.0001), region ($F_{9,144}$ = 50.11, $P$ < 0.0001), treatment × time ($F_{30,480}$ = 24.43, $P$ < 0.0001), treatment × region ($F_{18,288}$ = 18.11, $P$ < 0.0001), time × region ($F_{135,2160}$ = 39.21, $P$ < 0.0001), sex × time × region ($F_{135,2160}$ = 1.50, $P$ = 0.0003), treatment × time × region ($F_{270,4320}$ = 12.77, $P$ < 0.0001), sex × treatment × time × region ($F_{270,4320}$ = 1.20, $P$ = 0.015). Two-tailed paired $t$-test, VEH+KET vs NTX+KET, *P < 0.05, **P < 0.01.

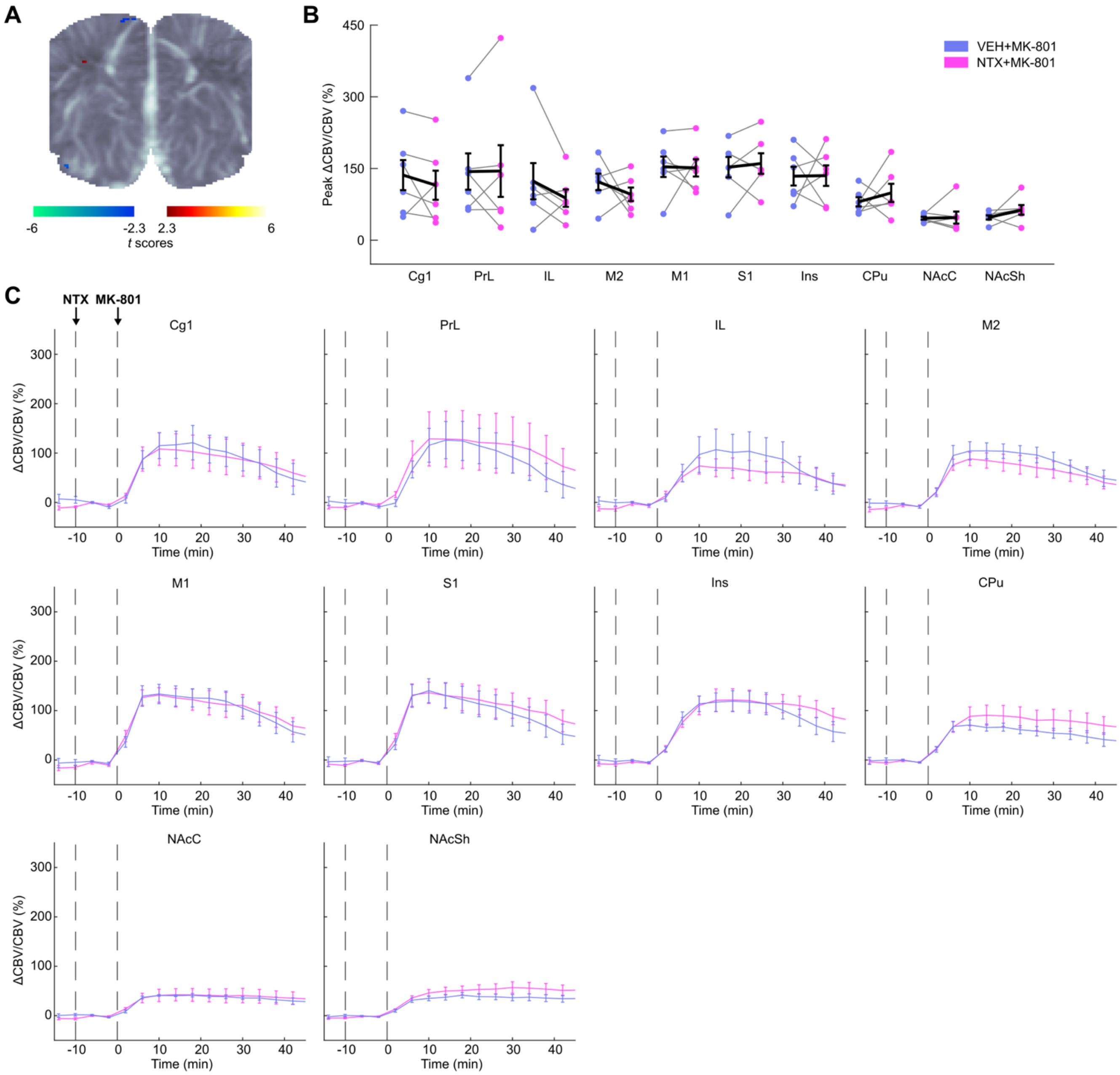


**Figure S4: Naltrexone pretreatment does not affect neural activity changes evoked by MK-801.** Male rats were administered either vehicle (VEH) naltrexone (NTX; 10 mg/kg) followed by an injection of MK-801 (0.1 mg/kg) after 10 min. The animals were imaged two times under the treatment conditions of VEH+MK-801 and NTX+MK-801. (**A**) Functional map at bregma +2.5 mm. The t scores were calculated by contrasting the pixel-wise peak cerebral blood volume (CBV) in the two treatment groups. The t scores were thresholded to show only the statistically significant pixels (two-sided paired t-test, $P < 0.05$). This map was not corrected for multiple comparisons. n = 6 rats. (**B**) Peak cerebral blood volume (CBV) changes in individual rats. Region-wise two-sided paired t-tests revealed no significant effects. (**C**) Regional CBV signals in the segmented ROIs were averaged in 4-min intervals. Lines are mean and error bars are s.e.m. from n = 6 male rats. Three-way ANOVA; within-subjects factor of treatment, , brain region, and time. Significant effects: time ($F_{15,75}$ = 28.96, P < 0.0001), region ($F_{9,45}$ = 2.63, P = 0.015), time × region ($F_{135,675}$ = 3.48, P < 0.0001). Two-tailed paired t-tests revealed no significant time points.

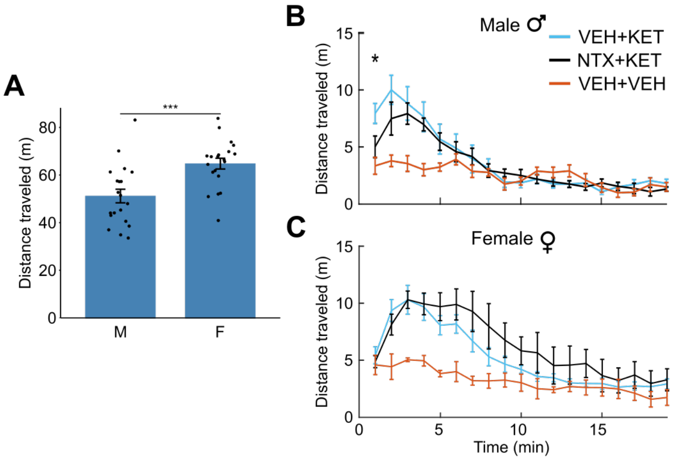


**Figure S5:** (**A**) Total distance traveled by male and female rats during habituation. Data are animal-wise average of two sessions. Two-tailed unpaired *t*-test, **P* < 0.0001. *n* = 20/group. (**B**, **C**) Distance traveled by male and female rats as a function of the time at day 1. Two-tailed unpaired *t*-test, VEH+KET vs NTX+KET groups, **P* < 0.05. *n* = 8/group in the VEH+KET and NTX+KET groups. *n* = 4 in the VEH+VEH group.

**Movie Captions:**

**Movie S1:** Brain-wide activity in two coronal planes following ketamine administration (10 mg/kg, intravenous). The cerebral blood volume (CBV) maps were calculated versus a pre-injection baseline and displayed overlaid on the respective power Doppler frames. Timestamp is in min:sec post ketamine.

**Movie S2:** Statistical maps comparing male rats administered ketamine (10 mg/kg, intravenous) and pretreated with the opioid receptor antagonist naltrexone (10 mg/kg, subcutaneous) (NTX+KET group) versus saline (VEH+KET group). The *t* scores for the two coronal planes are overlaid on the respective power Doppler frames. Timestamp is in min:sec post ketamine.
